# Disrupting Akt-Wnt/β-catenin signaling suppresses glioblastoma stem cell growth and tumor progression in immunocompetent mice

**DOI:** 10.1101/2025.07.25.666376

**Authors:** Md. Marzan Sarkar, Nathan Gonsalves, Amin Davarzani, Emma Mitchell, Amar M. Singh, Lohitash Karumbaiah, Steven L. Stice

## Abstract

Glioblastoma (GBM) is an aggressive primary malignant brain tumor in adults with a median patient survival of 12–18 months post-diagnosis. The PI3K/Akt and Wnt/β-catenin signaling pathways promote GBM cell growth, survival, invasiveness and therapeutic resistance. We hypothesize that inhibiting Akt and β-catenin, which are central regulatory proteins of the PI3K/Akt and Wnt/β-catenin pathway, will suppress GBM growth and progression. Our *in vitro* studies demonstrate that MK-2206, a pan-Akt inhibitor, effectively reduced cell viability, induced apoptosis, and inhibited β-catenin activity; consistently outperforming iCRT3, a β-catenin-TCF interaction inhibitor, in CT-2A mouse glioma cells, and N08-30 human glioma stem cells. Luciferase-expressing CT-2A cells were then intracranially implanted in C57BL/6J mice followed by MK-2206 treatment, and we observed a reduction in phosphorylated Akt and GSK-3β levels, consistent with disruption of the Akt and Wnt/β-catenin signaling axis causing tumor suppression. In summary, MK-2206 outperformed iCRT3 efficacy *in vitro*, and suppressed GBM progression, *in vivo*. These findings suggest that Akt inhibition via MK-2206 may offer a promising therapeutic strategy for treating GBM characterized by dysregulation of PI3K/Akt or Wnt/β-catenin pathways.

## 1. Introduction

Glioblastoma (GBM) is the most commonly occurring malignant brain tumor in adults [1]. Patients diagnosed with GBM have a low median rate of survival of 12–18 months, and standard-of-care (SOC) treatment consists of maximally safe surgical brain tumor resection and concurrent chemoradiotherapy [1-5]. A major drawback of the SOC chemotherapeutic agent temozolomide (TMZ) which is a DNA alkylating agent [6], is the development of chemoresistance observed in unmethylated tumors [7]. This is due to the enhanced activity of the DNA repair enzyme O^6^-methylguanine methyltransferase (MGMT), which accounts for approximately 50% of GBM patients [8-11]. The administration of TMZ chemotherapy in conjunction with low-intensity tumor treating fields (TTFields), or TMZ, in combination with anti-angiogenic agents such as bevacizumab, have been recently tested [5, 12-14]. However, these therapies have not yielded any significant survival benefits to GBM patients.

PI3K/Akt signaling pathway has been found to be hyperactive in GBM [15]. Studies have demonstrated hyperactivation of Akt in 50% of GBM cases, while mutations or deletions in phosphatase and tensin homolog (PTEN), a negative regulator of PI3K/Akt signaling, are found in 36–44% of GBM cases [3, 16-21]. Hyperactive PI3K/Akt pathway promotes cell survival and uncontrolled cell proliferation, apoptosis inhibition, drug resistance, immune evasion of glioma cells, and ultimately contributes to an immunosuppressive environment [22-24]. Furthermore, high levels of Wnt/β-catenin activity have also been observed in radio- and chemo-resistant GBM cells, and studies suggest that TMZ treatment activates the Wnt/β-catenin signaling pathway, resulting in TMZ- and radio-resistance in GBM [4]. Overall, these studies suggest that the Wnt/β-catenin signaling is strongly connected with poor clinical outcomes and a low patient-survival rate [25-27].

Akt hyperactivation occurs in most human cancers, including GBM, and can be triggered by Wnt activation of epidermal growth factor receptor (EGFR) signaling, resulting in PI3K activation, which promotes β-catenin–dependent tumor immune evasion [27, 28]. Hyperactive Akt phosphorylates and inhibits GSK-3β, which results in enhanced β-catenin accumulation and reciprocally, β-catenin/T□cell factor-4 (TCF4) interaction, directly regulates Akt1 in glioma [29, 30]. Furthermore, loss of APC in the β-catenin destruction complex promotes β-catenin accumulation and activates mTORC1 [31, 32] and mTORC1 promotes Akt activity [33, 34]. β-catenin, the major transcriptional driver of Wnt/β-catenin pathway, is also responsible for growth and maintenance of GBM cells through activation of Wnt target genes [35, 36]. These findings demonstrate that Akt and β-catenin play an important role in the crosstalk of PI3K/Akt and Wnt/β-catenin signaling pathways, facilitating convergent pathway signaling that promotes GBM cell proliferation, survival, and invasion. As GBM is characterized by abnormal activation of the PI3K/Akt and Wnt/β-catenin signaling pathways, blocking a common effector for both pathways, such as Akt or β-catenin or a dual-targeting strategy may prove to be more effective than conventional single pathway targeting therapies. Therefore, we hypothesize that inhibiting Akt and β-catenin, that are key central regulatory components of the PI3K/Akt and Wnt/β-catenin pathway, will suppress GBM growth and progression.

In our study, we assessed the inhibitory effects of the small molecule inhibitors MK-2206 2HCl (Akt and β-catenin inhibitor), and iCRT3 (β-catenin inhibitor), on the growth, proliferation, apoptosis, and β-catenin transcriptional activity of CT-2A mouse glioma cells and N08-30 human glioma stem cells, *in vitro*. MK-2206 is a potent and selective allosteric Akt inhibitor demonstrating *in vitro* and *in vivo* antitumor activity as a single agent [37] across cancer types, including solid tumors and has been demonstrated to cross the blood–brain barrier (BBB) [38]. iCRT3 disrupts the interaction between β-catenin and TCF of the Wnt signaling pathway [39]. Next, we evaluated the efficacy of MK-2206 in reducing the progression of aggressive, syngeneic CT-2A GBM tumors in immunocompetent C57BL/6J mice. Immunocompetent animal models provide a physiologically relevant tumor microenvironment with an intact host immune system [40]. CT-2A cells lack functional PTEN, which is associated with the activation of PI3K/Akt and subsequent activation of the Wnt/β-catenin pathway [29, 41]. CT-2A tumors parallel human high-grade gliomas (hHGG) both exhibit stem cell markers, high mitotic index and cell density, nuclear polymorphism, hemorrhagic pseudo palisading necrosis, and microvascular proliferations [42, 43]. Tumor-induced immunosuppression is also exacerbated by PTEN deficiency [3]. Therefore, an immunocompetent CT-2A model is well-suited to determine therapeutic effectiveness of interventions that suppress GBM progression caused by PTEN deficiency, and Akt and Wnt/β-catenin activation.

## 2. Materials and Methods

### 2.1. Cell Lines and Cell Culture

CT-2A mouse glioma cells (Millipore-Sigma, Cat. No.: SCC194) which harbor a PTEN mutation, were cultured in Dulbecco’s Modified Eagle’s Medium (DMEM), high glucose (Cytiva, Cat. No.: SH30081.02), supplemented with 10% fetal bovine serum (Neuromics, Cat. No.: FBS002.) and 1% penicillin/streptomycin (Gibco, Cat. No.: 15070-063).

Human glioblastoma stem cells (N08-30) were isolated from primary human GBM tissue, according to procedures approved by Emory University IRB protocol 45796 (D.J.B.) validated for glioblastoma stem cell markers [44-46]. N08 cells were maintained in Neurobasal-A Medium (Gibco, Cat. No.:10888-022) containing 1% penicillin-streptomycin and 0.5% L-glutamine (Gibco, Cat. No.: 35050061, supplemented with 1% N-2 supplement (Gibco, Cat. No.: 17502048), 2% B-27 supplement (Gibco, Cat. No.: 17504044), rhFGF (10ng/ml, R&D system Cat. No.: 233-FB-025/CR), hEGF (20ng/ml Gibco, Cat. No.: AF100151MG), and 0.4% heparin (STEMCELL, Cat. No.: 07980). Cells were incubated at 37°C with 5% CO_2_ and 95% relative humidity. Cells were fed with fresh medium, every two days, unless passaged, or harvested for *in vitro* and *in vivo* assays.

### 2.2. MTS Cell Viability Assay

CT-2A cells were seeded in 96-well plates at a density of 10K cells/well and left to attach for 24h in a 37°C cell culture incubator. Subsequently, cells were treated with or without drug/siRNA containing media and incubated for an additional 48h. Then 20μL MTS reagent (Abcam, Cat. No.: ab197010) was directly added to each well, and cells were incubated for 2-3 h. Absorbance at 490 nm was measured by using a microplate reader (SpectraMax iD5, Molecular Device). Absorbance values are directly proportional to the number of viable cells. Non-Linear Regression analysis was used to plot a dose-response inhibition curve (log-scale) and IC_50_ was calculated.

### 2.3. Detection of Cell Apoptosis

CT-2A cells were seeded on a 6-well plate at 600K per well and allowed to attach for ∼24h and then treated with or without drug media and incubated for another 24h. Apoptotic cells were determined using the Annexin V-FITC/PI Apoptosis Kit (Abnova, Cat.: KA3805). All procedures were performed according to the user manual with slight modification. Briefly, cells were washed with 4mL of cold PBS and resuspended in 100μL of 1X binding buffer. Then the binding buffer was removed from the cell pellet, and the cell pellet was resuspended again in cold 1X binding buffer. 5μL of Annexin V-FITC was added to 100μL cell suspension (120K cells) in each appropriate tube and incubated for 11 minutes at room temperature in the dark. Then 5μL of PI solution was added and incubated for 6 min at room temperature in dark. Finally, the binding buffer with Annexin V-FITC and PI was removed, and cells were resuspended in 100μL 1X binding buffer and analyzed by flow cytometry (Agilent Quanteon Flow Cytometer) immediately. A minimum of 13,000 single events/sample were recorded and gating thresholds were set from the vehicle group and uniformly applied across groups.

### 2.4. Caspase 3/7 Activity

Caspase-3/7 activity was measured by Caspase-Glo® 3/7 Assay System (Promega, Cat.: G8091) according to the manufacturer instructions with slight modifications. Briefly, cells were seeded in white-wall clear bottom 96-well plates at a density of 20K cells/well, allowed to attach for ∼24h and then treated with or without drug containing media (100μL) and incubated for another 6h. Cell culture plate was removed from incubator and allowed to equilibrate to room temperature for ∼10 min. Then 100µl of Caspase-Glo® 3/7 Reagent was added to each well, mixed well by pipetting and incubated for 1 hour at RT. Finally, luminescence was measured following treatment using the SpectraMax iD5 microplate reader. Here, luminescence is directly proportional to the caspase-3/7 activity.

### 2.5. Engineering CT-2A Mouse Glioma Cells

Pre-made lentivirus (GenTarget Inc, Cat. No.: LVP010-P-PBS) constitutively expressing Luciferase (Luc) reporter under the 8xTcf Wnt responsive promoter (mCMV) was used to infect CT-2A cells. This lentivirus also contains the puromycin selection marker, under the RSV promoter, and the cells were selected with 1µg/ml puromycin. This cell line was used for all subsequent *in vitro* experiments. CT-2A-Luc-Puro cells were generated by transducing cells with EF1-GFP-T2A-fLuc-Puro lentivirus (SignaGen Laboratories, Cat. No.: SL100329) at an MOI of 50. Cells were selected and maintained with 1µg/ml puromycin for >2 weeks to generate stable cell lines. This cell line was used for *in vivo* experiments.

### 2.6. Plasmid and siRNA Transfection

CT-2A cells were transfected using Lipofectamine RNAiMAX (ThermoFisher, Cat. No.: 13778030) by Akt siRNA (ON-TARGET plus siRNAs for Akt1, 2, 3 were purchased from Dharmacon). N08-30 cells were co-transfected using Lipofectamine 3000 (Invitrogen, Cat. No.:100022050) by 200 ng firefly luciferase expressing TOP and FOP-flash reporter plasmids (Millipore, MA, USA). Transfections were performed according to the manufacturer’s protocol. Briefly, cells were seeded in 24-well plate at 80K cells/well and allowed to attach. At 60-70% confluency, fresh media was added, and cells were transfected with siRNA/plasmid using transfection reagent. The concentrations of siRNAs or plasmids were chosen based on dose-response studies for knockdown efficiency.

### 2.7. TOPFlash/FOPFlash Reporter Assay

To evaluate the β-catenin transcriptional activity in human GBM cells, we used a pair of luciferase reporter constructs, TOPFlash and FOPFlash (a gift from Dr. Ken-ichi Takemaru, Stony Brook University). TOPFlash contains three copies of TCF/LEF binding site, and FOPFlash contains a mutated TCF/LEF binding site. Briefly, N08-30 cells were seeded in a precoated 24-well plate with Geltrex™ (Cat. No.: A1413202) at 80K cells/well and allowed to attach. At 60-70% confluence, fresh drug media was added, and cells were transfected with TOP and FOPFlash plasmid (200 ng) using Lipofectamine 3000 reagent (Invitrogen), according to the manufacturer’s protocol. After 48h of incubation, cells were collected and lysed with 1X passive lysis buffer (Promega, Cat. No.: E1941) using standard protocol conditions. From each treatment group, 10 µL cell lysate was directly added to each well of a 96-well white opaque plate. 50μL Luciferase Assay Reagent II (LARII, Promega, Cat. No.: E1500) was added, and luminescence signals were detected immediately using the microplate reader. Luciferase activity was normalized to total protein amount.

### 2.8. Luciferase Reporter Assay

To evaluate the β-catenin transcriptional activity in mouse glioma cells, we created a stable CT-2A cell line expressing luciferase signal as described in section 2.5. Briefly, CT-2A cells were seeded in a 24-well plate at 80K cells/well and allowed to attach. At 60-70% confluence, fresh drug media was added, or cells were transfected with Akt siRNA (100 nM) using Lipofectamine RNAiMAX reagent (Invitrogen, Cat. No.: 13778030) according to the manufacturer’s protocol. After 48h of incubation, cells were collected and lysed with 1X passive lysis buffer. From each treatment group, 10 µL cell lysate was directly added to each well of a 96-well white opaque plate. 50μL LARII was added, and luminescence signals were measured immediately using the microplate reader. Luciferase activity was normalized to total protein amount.

### 2.9. Western Blotting

Cells were washed with cold 1X PBS and lysed with RIPA buffer (Thermo Scientific, Waltham, Cat. No.: J63306-AK) supplemented with Halt Protease & Phosphatase Inhibitor cocktail (Thermo Scientific, Cat. No.:1861281) on ice for 30 minutes. Cell lysates were centrifuged at 12,000 RPM for 20 minutes at 4_C and supernatant was used to determine protein concentration by Pierce BCA protein assay kit (Thermo Fisher Scientific, Cat. No.: 23227). Proteins (20–30 μg) were separated through electrophoresis, using Mini-PROTEAN TGX Stain-Free Precast Gels (BIO-RAD, Cat. No.: 4568094) and then transferred to nitrocellulose membrane (Bio-Rad, Cat. No.: 1704271) using a turbo transfer system (BIO-RAD). The membranes were incubated with the respective antibodies from Cell Signaling Technology (p-AKT Ser473, Cat No.: 4060T; Akt pan, Cat No.: 4691T; p-GSK-3β Ser9, Cat No.: 5558T; GSK-3β, Cat No.: 12456T; and β-actin, Cat No.: 4970S) in 5% BSA in TBST at 4°C overnight, followed by incubation with secondary antibodies at room temperature for 1 h. Finally, protein bands were developed using Clarity western ECL substrate (BIO-RAD, Cat No.: 170-5060) using ChemiDoc MP imaging system (BIO-RAD). Protein amount was normalized to their respective β-actin loading control and band intensities were quantified by ImageJ software.

### 2.10. Quantitative Real-Time PCR

Total RNA was extracted from drug treated cells using Total RNA Kit (Cat. No.: R6834-01; Omega BIO-TEK, Norcross, GA, USA) according to the manufacturer’s instructions, and concentrations were measured using a Nano Drop 8000 (Thermo Scientific). cDNA was synthesized using iScript™ reverse transcription supermix (BIO-RAD; Cat.: 1708840). Primers were purchased from ThermoFisher and the assay ID of the primer sequences are: Myc; Mm00487804_m1, Ccnd1; Mm00432359_m1, Axin2; Mm00443610_m1 and GAPDH; Mm99999915_g1. qRT-PCR was performed in Quant Studio 5 Real Time PCR System (Applied Biosystems) using TaqMan™ Universal Master Mix II, no UNG (Fisher Sci. Cat. No.: 44-400-40) according to the manufacturer’s protocol. The 2^−ΔΔCT^ method was used to calculate the relative gene expression levels normalized to GAPDH and compared to control samples.

### 2.11. Animal Studies

All animal studies were approved by the University of Georgia Institutional Animal Care and Use Committee (IACUC), and all protocols were followed according to the National Institutes of Health’s Guide for the Care and Use of Laboratory Animals. Briefly, male C57BL/6J mice (6-8 weeks old, 18–30 g) were purchased from the Jackson Laboratory. The mice were housed in a temperature-controlled and light-controlled environment in the animal facility. Luciferase-expressing CT-2A cells (∼250K cells resuspended in 7μL DMEM medium) were injected in the right frontal cortex at coordinates +1.0 mm (AP), +1.5 mm (ML), and +3.0 mm (DV), relative to bregma, to induce mouse brain tumors. Bioluminescence imaging (BLI) was performed to confirm tumor uptake at Day-7, and mice were randomly divided into two groups (12 mice/group; MK-2206 and vehicle group). Intraperitoneal drug treatments were started on Day-8, according to the following regimen: MK-2206, 100 mg/kg body weight/day, and vehicle (20% β-captisol) at the same frequency until Day-28. Tumor growth and luciferase activity were monitored every 7 days using BLI imaging in NEWTON 7 (VILBER smart imaging). Animals were anesthetized using 1.5% isoflurane and, D-luciferin potassium salt (GOLDBIO, Cat. No.: LUCK-100) was injected (IP, 150mg/kg) 12-13 minutes prior to *in vivo* imaging for BLI imaging. Body weight and overall behavior were monitored daily for the first 7 days and then every other day until terminal euthanasia.

### 2.12. Brain Tissue Processing and Histology

At the 29-day time point, the mice were deeply anesthetized using 3-4% isoflurane gas and transcardially perfused with Phosphate Buffered Saline (PBS, pH 7.2) for 15-20 min to perfuse blood from vasculature in brain tissue. Post-perfusion, brains were carefully extracted, flash frozen on dry-ice, and stored at −80°C, until cryosectioning. The initial 0.5 mm from the start of the olfactory bulb (16 µm thick, 31 sections) was discarded, and total of 120 coronal sections (16 µm thick) were serially collected on 30 slides (4 sections per slide) along the rostro-caudal axis. The tissue sections were processed for hematoxylin and eosin (H&E) staining, and the stained slides were imaged at 20X magnification using a digital slide scanner (Aperio AT2) by the University of Georgia’s Histology Laboratory Core Facility.

### 2.13. Immunohistochemistry

One slide per animal (4 sections/slide) was used for immunostaining. Briefly, sections were fixed using 4% paraformaldehyde in 0.4 M sucrose for 20 minutes after rinsing with PBS and blocked using 5% BSA in PBST (0.5% Triton-X 100 in PBS) for 1 hour at room temperature (RT). Sections were then incubated with primary antibody (p-AKT Ser473, Cat No.: 4060T; p-GSK-3β Ser9, Cat No.: 5558; Cell Signaling Technology) overnight, at 4°C. Next day, the sections were rinsed in PBS, blocked again for 1 hour at RT, and incubated with Alexa Fluor 488 Goat anti-rabbit (Cat No.: A11008, Invitrogen) secondary antibody for 3 hours at RT. Finally, sections were stained with nuclear stain Hoechst 33342 (Invitrogen, Dilution 1:200 from 1mg/ml stock), airdried and cover slipped with Fluoromount-G mounting medium (Southern Biotech, AL) and stored at −20°C until imaged. The immunohistochemically stained brain sections were imaged at 20X on a Zeiss LSM880 with identical acquisition settings across groups; ROIs were segmented in CellProfiler [47] and signal was plotted as area normalized to Hoechst^+^ cells. A total of 5 mice from each treatment group were randomly selected for immunostaining studies.

## 3. Statistical analysis

GraphPad Prism 9.5.1 was used to perform all statistical analyses and data plotting. The data was tested for normality and equal variance assumptions using the Shapiro-Wilk test and F-test, respectively. Unless otherwise indicated, n refers to data from 3 biological experiments. Data indicated as Mean ± SD or Median ± Interquartile Range (IQR). Multi-group comparisons were performed using One-Way ANOVA with Tukey’s post hoc comparison. Unpaired two-tailed t-tests were used to compare data between two groups. Non-parametric statistics such as Mann-Whitney U-test was used for data that did not conform to the Gaussian distributions. For all tests, ^*^p<0.05 were considered statistically significant.

## 4. Results

### 4.1. Akt and β-catenin inhibitors promoted apoptosis, and suppressed the growth and proliferation of CT-2A cells, *in vitro*

To investigate the effect of Akt (MK-2206) and β-catenin (iCRT3) inhibitors on mouse glioma (CT-2A) cells (Figure **1A**), we performed the MTS cell viability assay. We found that MK-2206 and iCRT3 suppressed cell growth and proliferation in a concentration-dependent manner following 48h of treatment, when compared to vehicle group (Figure **1B-C**). The half-maximal inhibitory concentrations (IC_50_) values, for cell viability inhibition, were found to be 8.00 μM for MK-2206, and 40.86 μM for iCRT3.

**Figure 1:**
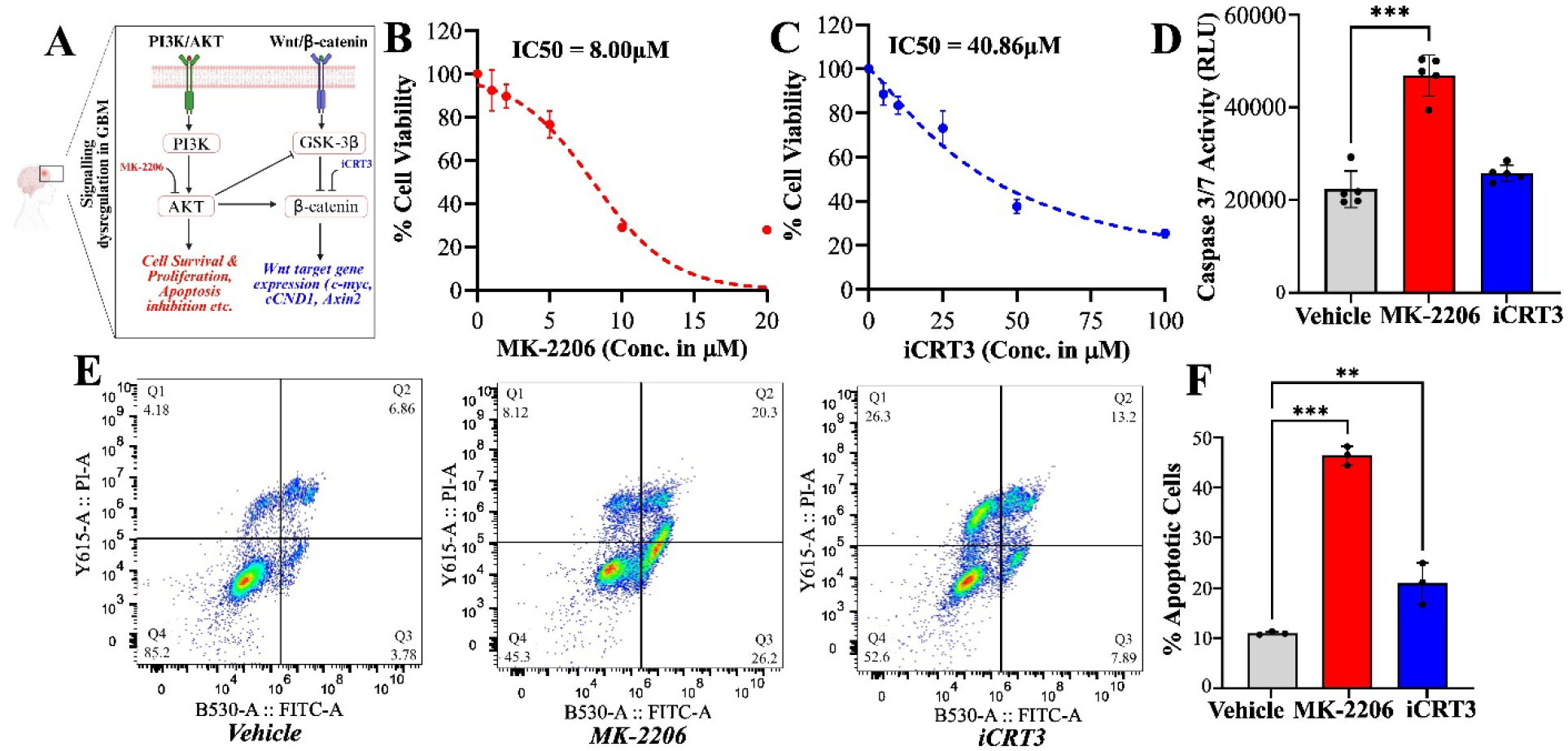
Akt and β-catenin inhibitors reduce CT-2A cell growth and induce apoptosis. **(A)** Schematic showing PI3K/AKT and Wnt/β-catenin pathway’s role in GBM cell growth and survival. **(B, C)** MK-2206 and iCRT3 dose-dependent changes in CT-2A cell growth and viability, respectively; (n=5) after 48h treatment. IC50 for MK-2206=8.0 μM and for iCRT3=40.86 μM. **(D)** Caspase 3/7 activity following 6h treatment of 10 μM MK-2206 or 50 μM iCRT3 (n= 5). (**E, F**) Flow cytometry analysis of CT-2A cells co-stained with propidium iodide (PI) and Annexin V following 24h of vehicle, MK-2206 (10 μM), or iCRT3 (50 μM) treatment (n=3). Data represents Mean ± SD from independent replicates. One-Way ANOVA with Tukey’s post hoc test; ^**^p<0.01 and ^***^p<0.001.

We measured Caspase-3/7 activity to determine whether MK-2206 and iCRT3 treatment led to increased apoptosis of CT-2A cells. We observed an increasing trend in Caspase-3/7 activity in iCRT3-treated cells following 6h treatment, while MK-2206 treatment significantly increased caspase activity, compared to the vehicle (Figure **1D**).

To confirm apoptosis induction in the inhibitor treated groups, the percentage of apoptotic cells were measured using Annexin V+ staining. Following 24h treatment, the percent apoptotic cells significantly increased in both, MK-2206 and iCRT3 treated cells, compared to vehicle (Figure **1E-F**).

Our data indicates that inhibition of Akt or β-catenin signaling decreases cell viability and induces apoptosis in CT-2A glioma cells. These findings suggest that Akt/β-catenin pathway contributes to cell survival and may represent a potential therapeutic target.

### 4.2. Wnt/β-catenin signaling is suppressed by MK-2206, but not iCRT3 treatment

We compared the effects of MK-2206 and iCRT3 on suppressing the Wnt/β-catenin signaling pathway. To determine whether MK-2206 and iCRT3 impacts β-catenin/TCF transcriptional -activity, we used luciferase expressing stable CT-2A cell lines having 8X TCF binding sites for β-catenin. When compared to vehicle, both MK-2206 and iCRT3 treated CT-2A cells (48h) exhibited significantly decreased luciferase reporter activity, indicating that Akt and β-catenin inhibitors downregulated β-catenin/TCF transcriptional activity in CT-2A cells (Figure **2A**).

**Figure 2:**
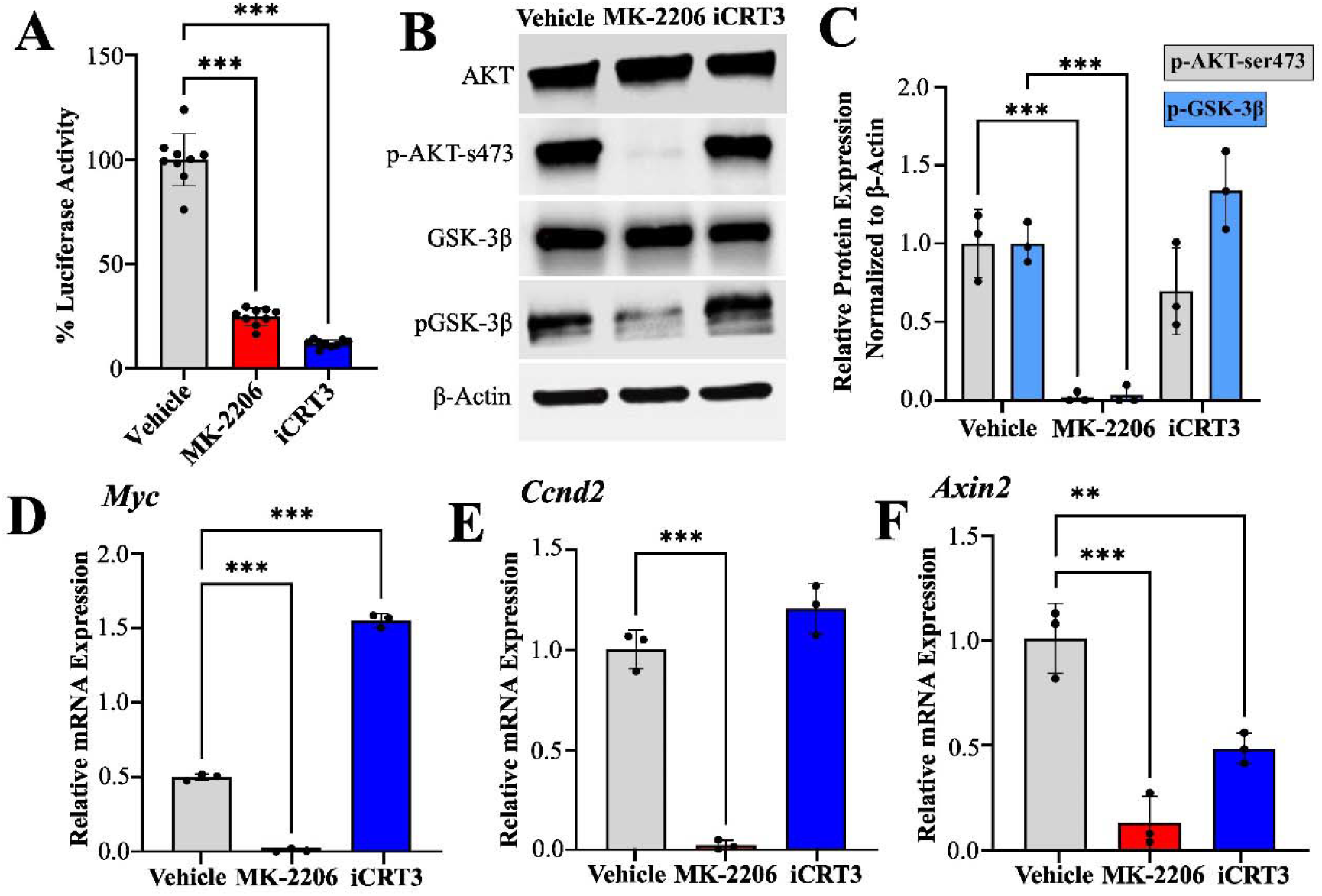
Akt and β-catenin inhibitors suppress transcriptional activity of β-catenin in CT-2A cells. **(A)** Percent luciferase activity of β-catenin using the luciferase reporter assay after Akt inhibitor MK-2206 (5μM) and β-catenin inhibitor iCRT3 (25μM) treatment of CT-2A cells for 48hr (n=9). **(B)** Immunoblotting of cell lysates from cells treated with MK-2206 (10 μM) and iCRT3 treatment (50 μM) for 1h compared to vehicle (n=3). **(C)** Densitometric analysis of Western blot bands for proteins from Fig 2C, using ImageJ. **(D-F)** qRT-PCR analysis of *Myc, Ccnd1*, and *Axin2* mRNA levels following 48-h treatment with MK-2206 (10 μM) and iCRT3 (50μM) (n=3, independent replicates). Data represents Mean ± SD. One-Way ANOVA with Tukey’s post hoc test; ^**^p<0.01 and ^***^p<0.001.

Western blot analysis was performed to investigate the effects of MK-2206 and iCRT3 treatment on phosphorylated Akt (p-AKT) and phosphorylated GSK-3β (p-GSK-3β) levels. CT-2A cells were treated with MK-2206 and iCRT3 for 1h, lysed and total protein was extracted for western blot analysis. MK-2206 treatment significantly down-regulated both p-AKT and p-GSK-3β proteins in CT-2A cells, compared to untreated vehicle (Figure **2B-C**). iCRT3 showed no significant effects on p-AKT or p-GSK-3β levels, which is in line with its mechanism of action of preventing TCF-β-catenin binding.

Given the positive correlation between hyperactive Akt and activation of Wnt/β-catenin signaling pathway, we hypothesized that the expression of known Wnt/β-catenin target genes *cyclin D1, c-Myc* and *Axin2* would be transcriptionally regulated by the Akt/β-catenin pathway in CT-2A cells. Results indicate that Akt inhibitor MK-2206 treatment significantly downregulated the expression of *cyclin D1, c-Myc* and *Axin2* (Figure **2D-F**). Though β-catenin inhibitor iCRT3 treatment significantly reduced the expression of *Axin2*, it concomitantly enhanced the expression of *c-Myc* and induced no significant changes in *cyclin D1*, when compared to vehicle, suggesting that MK-2206 can block Wnt/β-catenin signaling more effectively than iCRT3 (Figure **2D-F**).

These results suggest that Akt activation correlates with the levels of GSK-3β phosphorylation and Wnt/β-catenin target gene expression in GBM cells.

### 4.3. Akt and β-catenin inhibitors reduced human glioma stem cell growth and β-catenin transcriptional activity, while only Akt inhibition induced apoptosis

The effects of Akt and β-catenin inhibitors were then evaluated on N08-30, patient-derived glioma stem cell line, carrying both EGFR amplification and PTEN mutations. To investigate the inhibitory effects of MK-2206 and iCRT3 on N08-30, the MTS cell viability assay was performed. MK-2206 and iCRT3 suppressed glioma stem cell growth in a concentration-dependent manner following 48h of treatment compared to the vehicle group (Figure **3A-B**). IC_50_ values for inhibiting cell viability was found to be 8.17 μM for MK-2206, and >100 μM for iCRT3 (Figure **3A-B**). These results indicated that iCRT3 is less cytotoxic to patient-derived glioma stem cells, compared to MK-2206.

**Figure 3:**
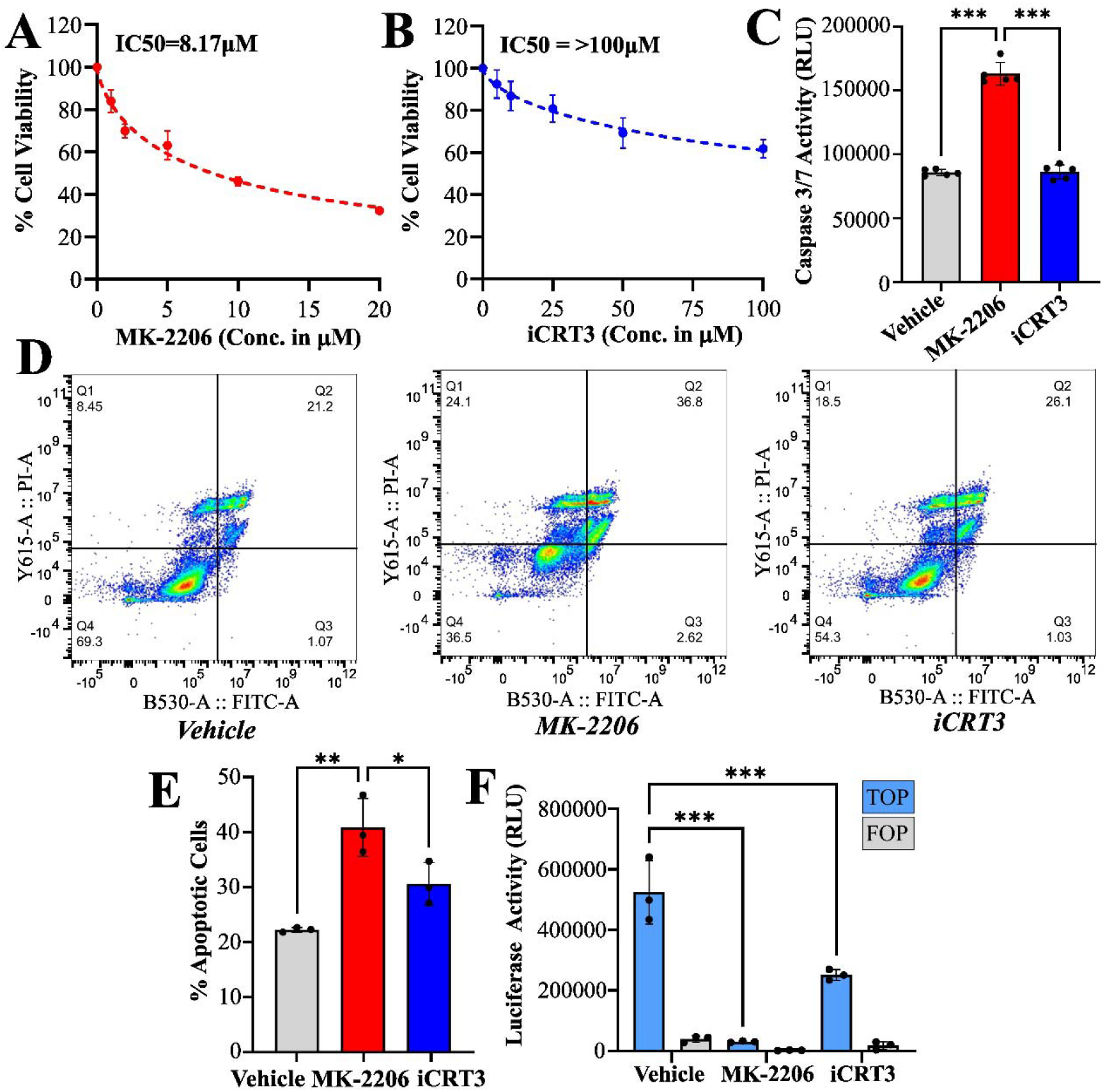
Akt inhibitor reduces N08-30 human glioma stem cell growth and induces apoptosis. **(A, B)** MK-2206 and iCRT3 dose-dependent changes in N08-30 cell growth and viability, respectively; after 48h treatment (n=5). IC50 for MK-2206=8.17 μM and for iCRT3=>100 μM. **(C)** Caspase 3/7 activity following 6h treatment of 10 μM MK-2206 or 50 μM iCRT3 (n= 5). **(D, E)** Flow cytometry analysis of N08-30 cells co-stained with propidium iodide (PI) and Annexin V following 24h of vehicle, MK-2206 (10 μM), or iCRT3 (50 μM) treatment (n=3). **(F)** β-catenin luciferase activity (RLU) of N08-30 cells treated with both Akt inhibitor MK-2206 (5μM) and β-catenin inhibitor iCRT3 (25μM) for 48h (n=3). Data represents Mean ± SD from independent replicates. One-Way ANOVA with Tukey’s post hoc test; ^*^p<0.05, ^**^p<0.01 and ^***^p<0.001.

The ability of MK-2206 and iCRT3 to induce glioma stem cell apoptosis was evaluated by quantifying caspase-3/7 activity and the percentage of Annexin V+ cells. Our results demonstrated that caspase-3/7 activity and percent apoptotic cells were significantly increased in MK-2206 treated cells, compared to vehicle and iCRT3 treatment (Figure **3C-E**). iCRT3 treatment neither enhanced caspase-3/7 activity, nor Annexin V+ cells, compared to the vehicle (Figure **3C-E**).

To determine whether MK-2206 and iCRT3 impacts β-catenin/TCF transcriptional activity in N08-30 human glioma stem cells, we transfected the cells with TOPflash and FOPflash reporter plasmids. When compared to the vehicle, MK-2206 and iCRT3 treated cells significantly decreased TOPflash reporter activity, indicating that both inhibitors downregulated β-catenin/TCF-induced transcription (Figure **3F**). There was no change in the FOPflash reporter activity, which served as mutant negative control.

Taken together, Akt inhibition by MK-2206 outperformed β-catenin inhibition by iCRT3 by suppressing patient-derived glioma stem cell growth, and inducing apoptosis *in vitro*, while both MK-2206 and iCRT3 significantly downregulated β-catenin/TCF-induced transcriptional activity in these cells.

### 4.4. Akt siRNA suppressed CT-2A cell growth, transcriptional activity of β-catenin, and reduced Wnt target gene expression

In our previous experiments, while both Akt inhibitor, MK-2206, and β-catenin inhibitor, iCRT3, showed promise, MK-2206 consistently outperformed iCRT3, prompting us to focus on Akt mediated Wnt/β-catenin pathway inhibition in future experiments. To confirm whether the inhibitory effect of MK-2206 was specifically mediated through Akt inhibition in our previous findings, we performed siRNA mediated knockdown of Akt in CT-2A cells.

Before assessing the functional impact from the loss of *Akt* isoforms (*Akt1, Akt2* and *Akt3*) on CT-2A cell growth, we first quantified their transcript levels (Figure **4A**). We observed that *Akt1* had the highest expression level, followed by *Akt2*, while *Akt3* exhibited the lowest levels (Figure **4A**). We then assessed the targeting of all Akt isoforms (*Akt1, Akt2, Akt3)* together. Subsequent viability assessments demonstrated that knockdown of all Akt isoforms together, resulted in a significant reduction in cell viability (Figure **4B**).

**Figure 4:**
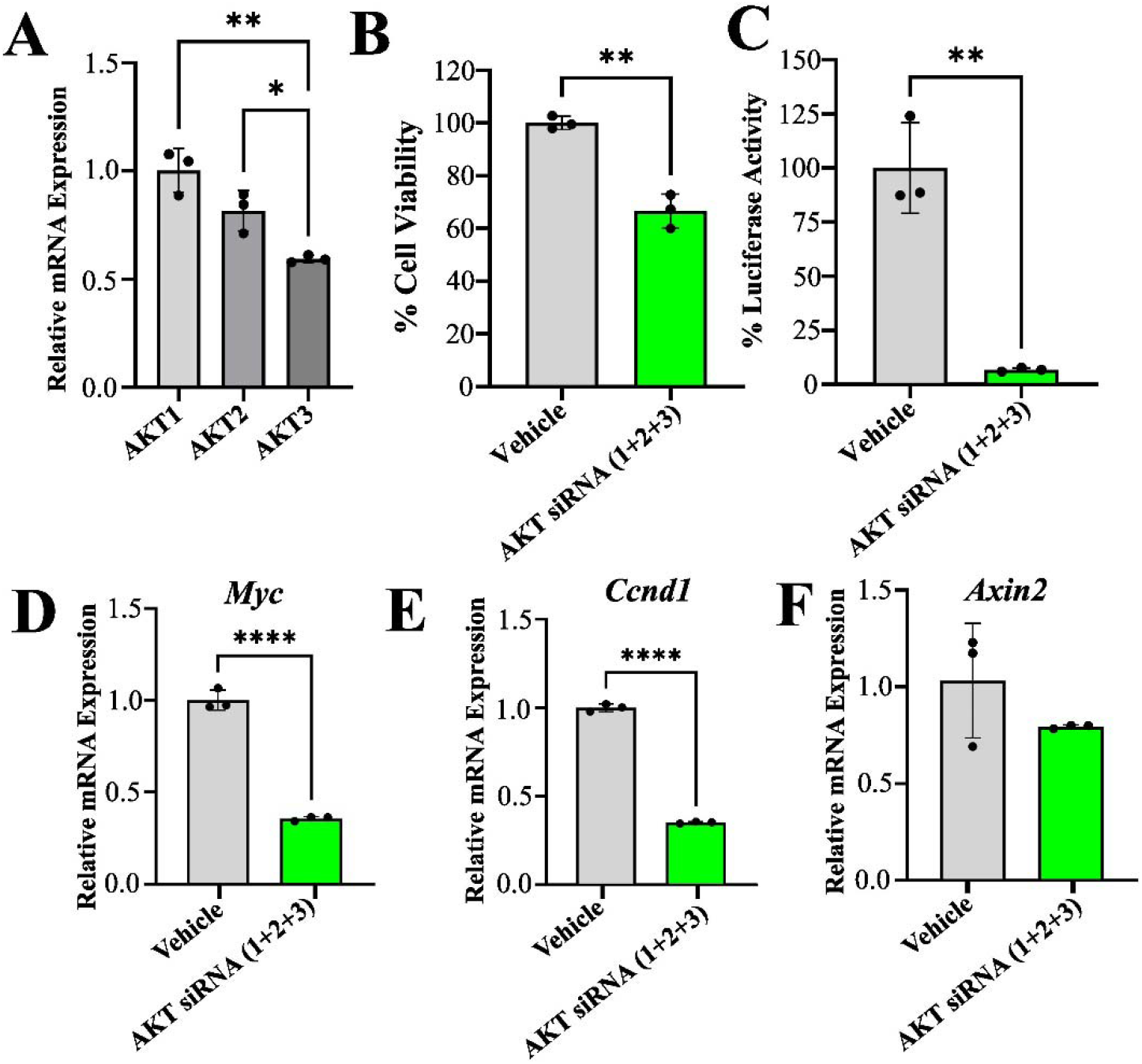
Akt siRNAs suppress CT-2A mouse glioma cell growth and Wnt/β-catenin signaling. (**A**) Relative mRNA expression of Akt isoforms (*Akt1, Akt2* and *Akt3*) in CT-2A cells. (**B**) Percent cell viability following 48h treatment by combinatorial Akt siRNA (1+2+3) in CT-2A cells, compared to the vehicle (n= 3). (**C**) Percent luciferase activity of β-catenin using Akt siRNA (1+2+3) following 48h treatment in CT-2A cells, compared to the vehicle (n=3). (**D-F**) qRT-PCR analysis of *c-Myc, Ccnd1*, and *Axin2* mRNA levels following 48h treatment with Akt siRNA, compared to the vehicle (n=3). Data represents Mean ± SD from independent replicates. One-Way ANOVA with Tukey’s post hoc test; ^*^p<0.05, ^**^p<0.01 and ^****^p<0.0001.

To determine whether knocking down of all Akt isoforms affects β-catenin/TCF-interaction and Wnt/β-catenin target gene expression, we performed luciferase reporter assay and qRT-PCR on CT-2A cells. Knockdown of all *Akt* isoforms led to a significant decrease in β-catenin/TCF-mediated transcription, as observed through a reduction in luciferase activity (Figure **4C**). Furthermore, simultaneous knockdown of all three Akt isoforms resulted in a significant reduction in *cyclin D1* and *Myc* (Figure **4D-E**) while *Axin2* level showed a decreasing trend, compared to the vehicle (**4F**).

These findings verify that the observed effects of MK-2206 resulted from specific AKT inhibition, not non-specific activity.

### 4.5. MK-2206 suppressed glioma tumor progression in immunocompetent mice

From our findings on mouse glioma and human glioma stem cells, the Akt inhibitor MK-2206 consistently outperformed iCRT3, and therefore we limited our investigations to studying Akt mediated Wnt/β-catenin pathway inhibition in suppressing mouse glioma cell growth, *in vivo*. To evaluate the inhibitory effect of MK-2206 on tumor progression *in vivo*, we performed orthotopic inoculations of luciferase expressing mouse CT-2A cells into immunocompetent C57BL/6J mice. CT-2A-Luciferase (CT-2A-Luc) cells were intracranially inoculated in the right frontal cortex of C57BL/6J mice. Tumor uptake was confirmed at Day-7 following an intraperitoneal (IP) D-luciferin injection (150mg/kg body weight) and subsequent bioluminescence (BLI) imaging. Mice were then randomly divided into two groups with one group receiving daily IP injections of the MK-2206 (100mg/kg body weight) and other group receiving 20% β-captisol as vehicle treatment (Figure **5A**).

**Figure 5:**
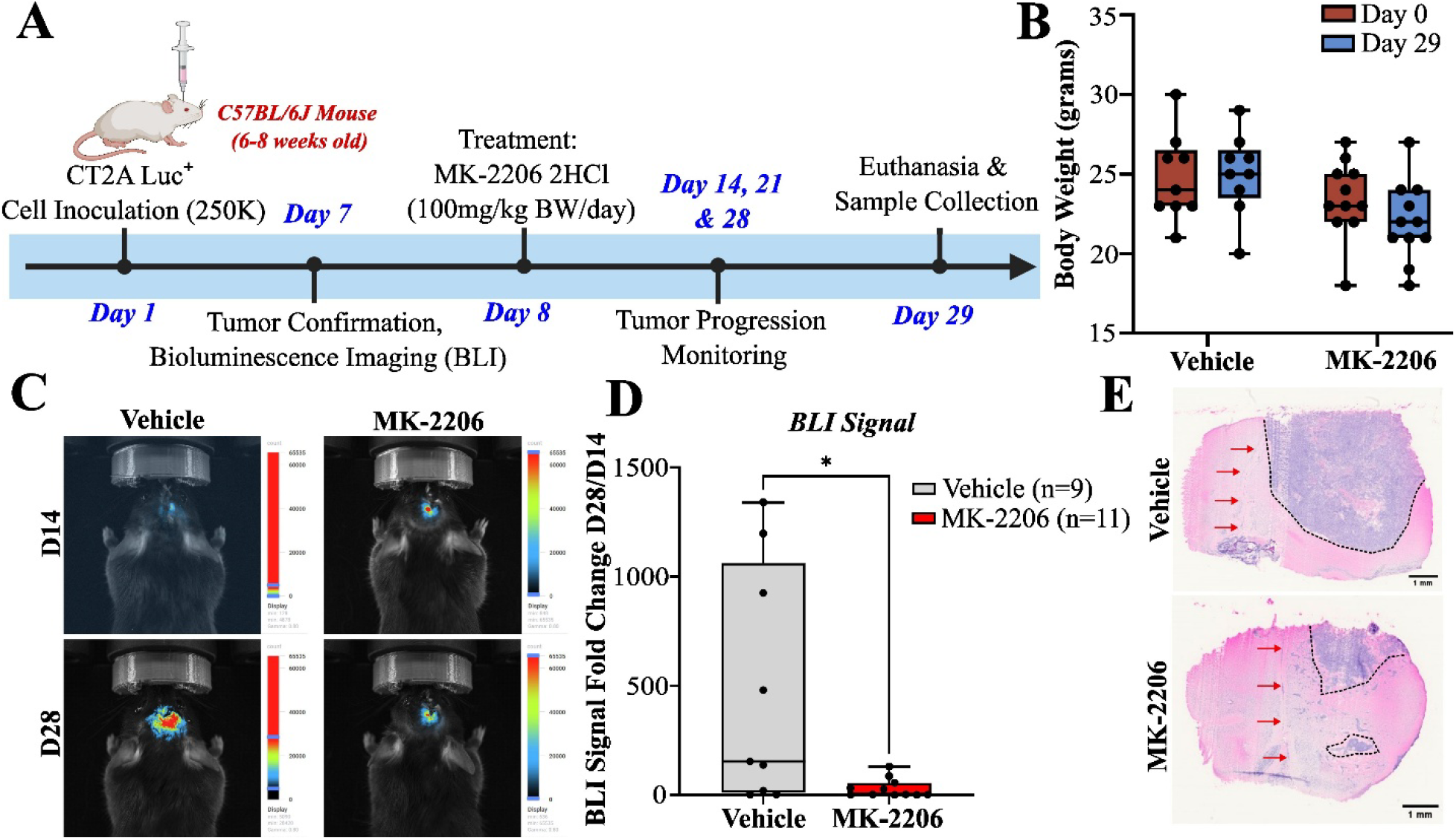
Akt inhibition significantly suppresses GBM tumor progression, *in vivo*. **(A)** Timeline of CT-2A syngeneic GBM induction in C57BL/6J mice, followed by MK-2206 treatment, with BLI imaging, terminal euthanasia and brain tissue collection. **(B)** Body weight (in grams) of mice at Day 0 and Day 29 after intraperitoneal administration of MK-2206 (vehicle, n=9 and MK-2206, n=11). **(C)** Representative images of vehicle and MK-2206 treated mice undergoing BLI imaging. **(D)** Box plot showing BLI signal fold change (Day 28/Day 14) for vehicle (n=9) vs MK-2206 (n=11) treated groups. **(E)** Representative hematoxylin and eosin (H&E) staining with tumor bulk from vehicle and MK-2206 treatment group. Red arrow indicates the midline, and dotted black border indicates hyper-nucleated area. Scale bar – 1mm. Data represents Median ± IQR. Two-Way Repeated Measures ANOVA or Two-tailed unpaired Mann-Whitney U-test. ^*^p<0.05.

Mice body weights and any abnormalities (e.g., coordination, tremor, head tilting, eye position, ruffled fur) were monitored throughout the study period (data not shown). We observed a decreasing trend in mice body weight from Day-0 to Day-29 in the MK-2206-treated animals. However, the observed weight loss was not found to be statistically significant (Figure **5B**).

Tumor progression was monitored over time using BLI imaging (Figure **5C**). Our results demonstrated that the tumor size (BLI signal at Day-28, compared to Day-14) of the MK-2206-treated group was significantly reduced, compared to vehicle group (Figure **5D**). At study endpoint of 29-days, mice were euthanized, and tumor tissues were collected for histological analysis. Representative hematoxylin and eosin (H&E) staining confirmed the tumor burden reduction in MK-2206 treated group compared to vehicle group (Figure **5E**), which was consistent with the BLI results. Three mice were excluded for lack of tumor uptake confirmed by BLI at Day 7, and one vehicle mouse died during study, finally MK-2206 (n=11) and vehicle (n=9) were used for analysis.

### 4.6. Akt inhibition suppressed downstream protein phosphorylation in tumor tissue

To determine whether intraperitoneal MK-2206 administration inhibited the expression of the Akt and Wnt/β-catenin signaling pathway, tumor tissue sections were analyzed for p-AKT and p-GSK3β expression through immunohistochemical analyses. The levels of p-AKT Ser473 and p-GSK-3β Ser9 were significantly downregulated in MK-2206 treated mice, compared to vehicle group, indicating that MK-2206 treatment suppressed Akt activity in GBM tumors, *in vivo* (Figure **6A-D**).

**Figure 6:**
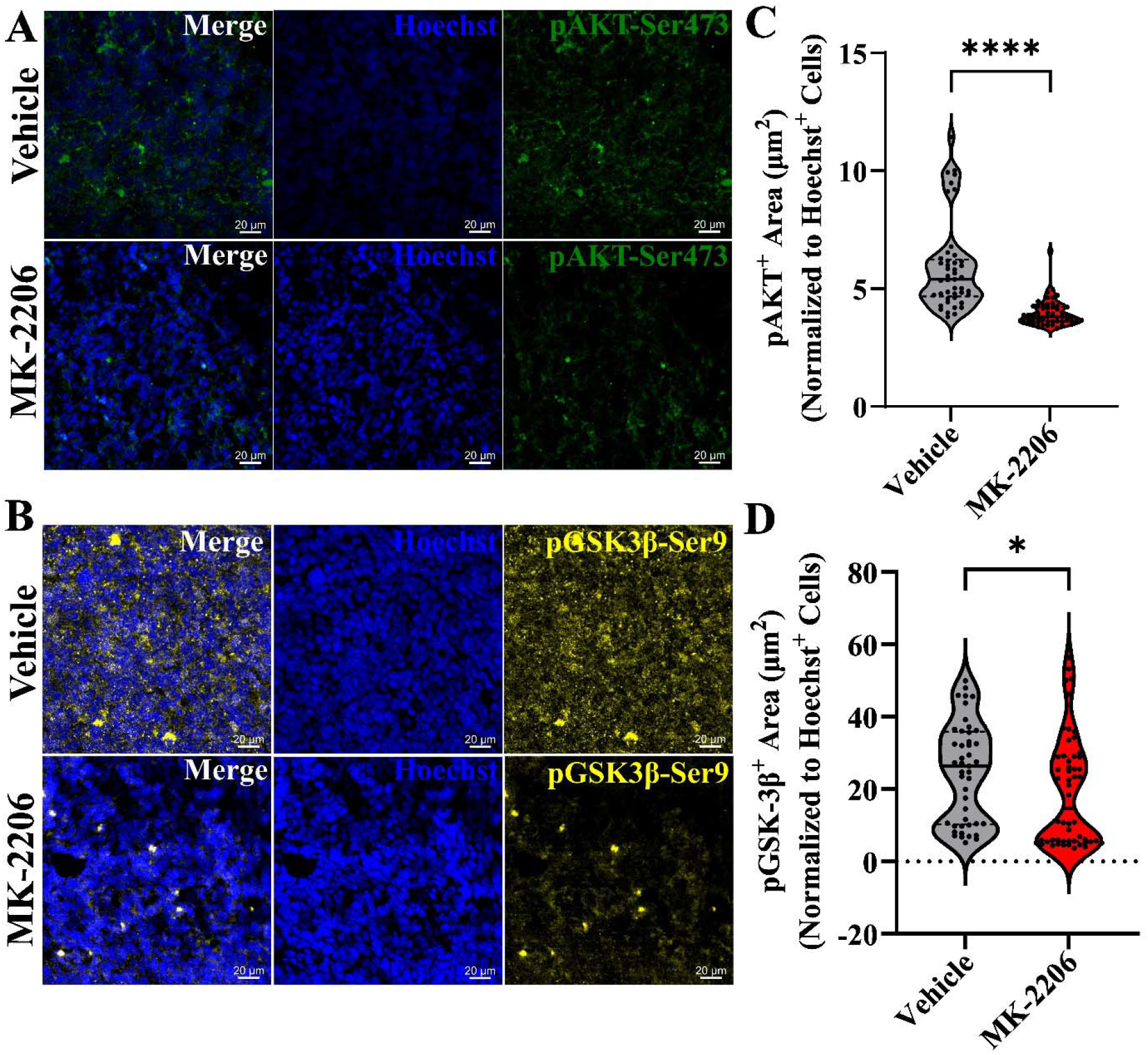
MK-2206 treatment suppresses p-AKT and p-GSK-3β expression in tumor tissues. Representative IHC images showing p-AKT Ser473 (**6A)** and p-GSK-3β Ser9 **(6B)** expression in MK-2206 treated mice compared to vehicle group. Hoechst –blue, pAKT-Ser473 – green and p-GSK-3β-Ser9 – yellow. Violin plots quantifying area of p-AKT Ser473 (**6C)** and p-GSK-3β Ser9 (**6D)** across vehicle and MK-2206 treated mice. Each data point (n) represents one ROI obtained from three sections per mouse and a total of 5 mice per group. Scale Bar - 20µm. Data represents Median ± IQR. Two-tailed Mann-Whitney U-test. ^*^p<0.05 and ^****^p<0.0001.

Altogether, our *in vivo* data demonstrated that Akt inhibition by MK-2206 significantly suppressed glioma tumor progression, likely by inhibiting both PI3K/Akt and Wnt/β-catenin signaling. This supports the notion that Akt inhibitors may have therapeutic utility for GBM with elevated PI3K/Akt and Wnt/β-catenin signaling.

## 5. Discussion

PI3K/Akt and Wnt/β-catenin signaling are two key drivers of GBM malignancy and therapeutic resistance [20, 27, 29, 48]. Genetic alterations in the PI3K/Akt/mTOR signaling cascade, a key driver of cell survival and apoptosis inhibition, were detected in 88% of all GBM cases [49]. While expression of multiple proteins in the pathway are dysregulated, hyperactivation of Akt alone occurs in approximately 50% of GBM cases [16, 17, 22]. Additionally, β-catenin, the major transcriptional driver of Wnt/β-catenin pathway is considered a prognostic marker, due to enhanced mRNA and protein expression in GBM [50, 51]. The crosstalk between these two pathways contributes to GBM’s invasive phenotype. Akt-mediated Wnt/β-catenin activation drives the transcriptional activation of Wnt target genes such as *Myc, cyclin D1*, and *Axin2*, which are responsible for GBM growth, proliferation, and immune evasion [27, 36, 48, 52]. However, the role of Akt mediated Wnt/β-catenin in glioma stem cell growth and proliferation has not been sufficiently elucidated. To address this knowledge gap, we evaluated the therapeutic potential of inhibiting Akt and β-catenin signaling in mouse and human patient-derived glioma stem cells, *in vitro*, and validated findings using a syngeneic, immunocompetent mouse GBM model, *in vivo*.

We compared the inhibitory effects of MK-2206, a potent and selective allosteric Akt inhibitor [37]; and iCRT3, a β-catenin-TCF interaction inhibitor [53], in both, mouse glioma cells (CT-2A), and patient-derived glioma stem cells (N08-30). The enhanced apoptotic response and reduced cell viability observed in MK-2206 treated group demonstrates the central role of Akt in regulating pro-survival pathways in GBM. Mechanistically, Akt inhibition promotes apoptosis by reducing the phosphorylation of pro-apoptotic proteins such as BAD and BIM [54, 55] and by activating apoptotic gene expression via FOXO3a dephosphorylation [56]. Additionally, Akt inhibition suppresses NF-κB activity and reduces the expression of anti-apoptotic targets like *Bcl-xL* [57, 58]. These mechanisms highlight MK-2206’s pro-apoptotic efficacy over iCRT3 in GBM models.

Overexpression of *c-Myc* and *cyclin D1*, the downstream targets of Akt and Wnt/β-catenin signaling contributes to uncontrolled cell proliferation and inhibition of apoptosis in gliomas [59-62]. p-Akt promotes cell survival and proliferation by modulating downstream targets including mTOR, FOXO3a and MDM2, while simultaneously inhibiting GSK-3β phosphorylation [63]. GSK-3β inhibition stabilizes β-catenin and increases the expression of oncogenic Wnt/β-catenin target genes such as *c-Myc* and *cyclin D1* [64, 65]. In our findings, MK-2206 downregulated p-Akt and p-GSK-3β in mouse glioma cells, but iCRT3 lacked this effect, indicating limited target modulation. Similar to our results, previous studies using MK-2206 as single agent or in combination have shown that Akt inhibition leads to decreased phosphorylation of Akt, contributing to reduced GBM cell proliferation and enhanced apoptosis [66, 67]. The unexpected upregulation of *c-Myc*, by β-catenin inhibitor iCRT3 might be due to the compensatory effects of other signaling cascades [68]. These findings emphasize Akt’s role as a central regulator linking PI3K/Akt and Wnt/β-catenin signaling in GBM.

Further evidence for therapeutic Akt inhibition is supported by the recent (2023) FDA-approval of the pan-Akt inhibitor Capivasertib (AZD5363) [69, 70]. Both MK-2206 and Capivasertib act as pan-Akt inhibitor [67, 71, 72] and suppress cell growth and proliferation in a range of cancers through targeting downstream signaling pathways including mTORC1, GSK-3β, and FOXO3a [67, 73, 74]. In preclinical and Phase I clinical studies, Capivasertib has demonstrated radio-sensitizing effects and reduced cell viability targeting Akt and GSK-3β phosphorylation in oral and breast cancer model [75, 76] indicating a mechanistic profile overlapping with MK-2206 [72]. Results from these studies have demonstrated promising anti-tumor activity across multiple cancers by suppressing Akt signaling [77, 78]. Our findings with MK-2206 similarly support the therapeutic value of targeting Akt, in PTEN-deficient Akt-driven glioblastoma.

From our *in vitro* studies, the Akt inhibitor MK-2206 consistently outperformed iCRT3, and therefore we limited our investigations to studying Akt mediated Wnt/β-catenin pathway inhibition in suppressing GBM cell growth, *in vivo*. Previous studies investigating the therapeutic effects of the Akt inhibitor MK-2206 in GBM models were often limited by the use of animals with low tumor burden or immunodeficient backgrounds, limiting the clinical relevance of tumor-host interactions [27, 38, 67, 79]. Consequently, the single-agent therapeutic potential of MK-2206 against highly invasive GBM tumors in immunocompetent animals has not yet been fully investigated. In our study, we employed a syngeneic, immunocompetent C57BL/6J mouse model with established intracranial CT-2A tumors and monitored treatment effects using BLI imaging. MK-2206 treatment significantly suppressed tumor progression compared to vehicle group, which was consistent with previously reported findings in immunodeficient mouse models [67]. H&E staining of MK-2206-treated tumor tissues further supported tumor suppression revealing reduced hyper-nucleated and necrotic area, and a reduced midline shift, which are distinguishing features of aggressive and invasive GBM phenotypes [80, 81]. Although MK-2206 treatment markedly decreased tumor burden, the residual tumors displayed less defined tumor borders and local infiltration. This could be due to the highly invasive nature of glioblastoma, as eradication of all tumor cells is rarely achieved even in aggressive preclinical models. Additionally, decreased phosphorylation of Akt and GSK-3β in tumor tissue, confirming GSK-3β–mediated Wnt/β-catenin signaling inhibition by Akt inhibitor MK-2206.

Overall, our *in vitro* and *in vivo* findings strongly support the therapeutic potential of targeting hyperactive Akt with MK-2206 in GBM driven by aberrant Akt-Wnt/β-catenin axis. Future studies should evaluate MK-2206 in combination with potential drugs or chemoradiotherapy (or radiotherapy) agents including brain pharmacokinetic/pharmacodynamic (PK/PD) profiling and biomarker stratification to enhance translational potential for GBM treatment.

## Data Availability Statement

All data needed to evaluate the conclusions in the paper are present in the paper itself. Data can be obtained on demand by contacting the corresponding author: (S.L. Stice) and (L. Karumbaiah).

## Acknowledgement and Funding

We thank UGA’s Biomedical Core Facility (BMC) for providing us with access to confocal microscopes and *in vivo* Bioluminescence imaging facilities. We thank Shanmathi Ramasubramanian for assisting during pilot *in vivo* studies. We also thank Dr. Ken-ichi Takemaru, Stony Brook University, for providing luciferase reporter plasmids. This research was supported by the Georgia Research Alliance awarded to Steven Stice.

## Author Contributions

MMS, AS, LK and SLS contributed to the study conception and design; MMS performed all *in vitro* experiments; AS and MMS created luciferase expressing CT-2A cell lines; MMS, NG and AD performed *in vivo* tumor inoculation study; MMS conducted all *in vivo* imaging and analysis; MMS and NG performed immunohistochemical staining and imaging, and AD performed IHC analysis; EM assisted with cell culture and caspase 3/7 assay; MMS wrote the original draft and, AS, NG, AD, LK and SLS contributed to writing and revision of the manuscript.

The manuscript was written through contributions of all authors, and all authors have provided approval to the final version of the manuscript.

